# Trioxane-based MS-cleavable Cross-linking Mass Spectrometry for Profiling Multimeric Interactions of Cellular Networks

**DOI:** 10.1101/2024.08.06.606913

**Authors:** Clinton Yu, Eric Novitsky, Xiaorong Wang, Ignacia Echeverria, Scott Rychnovsky, Lan Huang

## Abstract

Cross-linking mass spectrometry (XL-MS) is a powerful technology for mapping protein-protein interactions (PPIs) at the systems-level. By covalently connecting pairs of proximal residues, cross-linking reagents provide distance restraints to infer protein conformations and interaction interfaces. While binary cross-links have been remarkably informative, multimeric cross-links can offer enhanced spatial resolution to facilitate the characterization of dynamic and heterogeneous protein complexes. However, the identification of multimeric cross-links remains extremely challenging due to fragmentation complexity and the vast expansion of database search space. Here, we present a novel trioxane-based MS-cleavable homotrifunctional cross-linker TSTO, which can target three proximal lysine residues simultaneously. Owing to its unique structure and MS-cleavability, TSTO enables fast and unambiguous identification of cross-linked peptides using LC-MS^n^ analysis. Importantly, we have demonstrated that the TSTO-based XL-MS platform is effective for mapping PPIs of protein complexes and cellular networks. The trimeric interactions captured by TSTO have uncovered new structural details that cannot be easily revealed by existing reagents, allowing in-depth description of PPIs to facilitate structural modeling. This development not only advances XL-MS technologies for global PPI profiling from living cells, but also offers a new direction for creating multifunctional MS-cleavable cross-linkers to further push structural systems biology forward in the future.

## INTRODUCTION

Cross-linking mass spectrometry (XL-MS) has emerged as a transformative technology for interactomics and structural proteomics^1–4^. By covalently linking proximal amino acid residues within or between proteins, XL-MS provides unique insights into the architecture of protein complexes and the spatial arrangement of individual protein domains. Due to various innovations, this methodology has revolutionized structural biology by complementing traditional structural techniques and offering a more holistic view of protein assemblies in their native environments ^4–8^. Over the years, XL-MS has significantly evolved with the incorporation of various functionalities and targeting groups in cross-linker design, broadening its applications and enhancing the scope of XL-MS studies ^9–13^. In particular, diverse MS-cleavable cross-linking reagents have been developed to facilitate the detection and identification of cross-linked peptides ^1, 3–4, 14–15^, enabling effective proteome-wide analyses of cellular networks to elucidate their structural organization from intact cells, subcellular organelles, tissues and clinical samples.

Up to now, the majority of cross-linking reagents are bifunctional (carrying two reactive groups) and current XL-MS analyses have focused on the identification of cross-links between two residues to infer pair-wise interactions. While binary cross-links have been shown to be immensely effective in PPI mapping, multimeric cross-links can offer enhanced spatial resolution and provide additional distance restraints to facilitate structural analysis of protein complexes. Furthermore, these higher-order cross-links can uncover novel interaction sites and conformations that are not easily detectable with binary cross-linkers. The resulting multimeric interactions can lead to the discovery of new functional insights and regulatory mechanisms within protein complexes—especially for dynamic and heterogeneous ones that contain subcomplexes and multivalent interactions. Although multimeric cross-links can be formed by bifunctional cross-linkers, the difficulty of their identification has rendered them largely invisible. Due to the additional expansion of search space associated with each successive peptide, traditional non-cleavable cross-linking reagents are incapable of this task. As a result, MS-cleavable cross-linking reagents are critical for alleviating this issue by physically separating cross-linked constituents to facilitate their individual identifications^1, 14–15^. During the course of our study, a homotetrafunctional MS-cleavable cross-linking reagent utilizing four NHS ester-targeting groups (aka Bisby) was very recently reported ^16^. However, the additional bonds required for cross-linking each successive peptide complicate cross-link fragmentation and identification. Therefore, new approaches are needed to permit effective identification of multimeric cross-links, especially for proteome-wide PPI profiling.

To this end, we aimed to design a homotrifuctional MS-cleavable cross-linker that simultaneously targets three residues, but more importantly streamlines the peptide identification process by utilizing a core structure that enables the synchronous release of all cross-linked peptides in a single step. This will allow us to expand the detection of protein connectivity and provide additional restraint information for improved structural elucidation at the systems-level. Here, we present the design, synthesis and characterization of a novel membrane-permeable, MS-cleavable trifunctional cross-linker, TSTO (tris-<u>s</u>uccinimidyl tri<u>o</u>xane) to enable simultaneous capture and identification of trimeric PPIs. XL-MS analysis of human 26S proteasomes has demonstrated that TSTO is effective in cross-linking protein complexes and that the resulting cross-linked peptides display unique and predicable fragmentation during collision-induced dissociation (CID), enabling their simplified and accurate identification using multistage mass spectrometry (MS^n^). Importantly, we have captured novel trimeric interactions to better define interfaces between proteasome subcomplexes. Moreover, TSTO is membrane permeable and has been successfully applied for *in vivo* cross-linking of HEK293 cells, revealing an XL-proteome containing 1512 proteins and 1242 interactions, expanding PPI coverages of cellular networks. Apart from binary interactions, the capture of trimeric interactions has facilitated the detection and AlphaFold-based modeling of protein oligomers. In summary, we show that TSTO can uncover novel interaction sites and conformations, leading to more detailed PPI networks and advancing our understanding of cellular processes and biological function.

## RESULTS

### Developing a Novel Trioxane-based MS-cleavable Homotrifunctional Cross-linker TSTO

In order to capture trimeric interactions, we designed a novel trioxane-based cross-linker TSTO and accomplished its synthesis through five steps (Figure 1A). As shown, TSTO carries a unique symmetrical structure comprising three NHS esters connected via a central trioxane group ^17–18^ (Figure 1A), permitting concurrent cross-linking between three lysine residues to form a trimeric cross-link among three individual peptides (aka tripeptide tri-link, [α,β,γ]) (Figure 1B, Type I). In comparison to a traditional cross-link between two individual peptides, accurate identification of a tri-link would be much more challenging due to further expansion of database search space (n^3^). Therefore, the design of the central trioxane is crucial as it carries three equal MS-cleavable bonds that are weaker than peptide bonds that can be concurrently cleaved using collision-induced dissociation (CID) to simultaneously release all three cross-linker arms in a single step. This leads to physical separation of the three cross-linked peptide constituents, yielding three fragment ions during MS^2^ analysis that can be subjected to MS^3^ for sequencing (Figure 1B, Type I). As shown, trioxane cleavage results in an identical and defined aldehyde remnant (AR) on each peptide constituent, allowing for their unambiguous identification. In addition, this minimizes the total number of MS^2^ fragments, simplifying ion selection for subsequent MS^3^ analysis. In addition to tripeptide TSTO cross-links, tri-links can be formed between two peptides, linking one lysine in one peptide (α) and two lysines in another peptide (β) (aka dipeptide tri-link, [α-β_2_]) (Figure 1B, Type II). For dipeptide tri-links, two fragment ions would be observed in MS^2^, one corresponding to a peptide carrying a single AR-modified lysine and the other representing a peptide carrying two AR-modified lysines. Finally, TSTO cross-linking can yield traditional cross-links in which two lysines from two different peptides are cross-linked while the third NHS ester of TSTO is hydrolyzed (aka, dipeptide bi-link, [α-β]) (Figure 1B, Type III). MS^2^ analysis of a dipeptide bi-link would yield two fragment ions corresponding to peptides each carrying a single AR-modified lysine, while the hydrolyzed arm is released as a neutral loss. These three types of TSTO cross-links are the most structurally informative products for PPI mapping and can be identified using the same LC-MS^n^ workflow that has been previously established for our sulfoxide-containing MS-cleavable cross-linkers ^1, 4, 19^.

**Figure 1.**
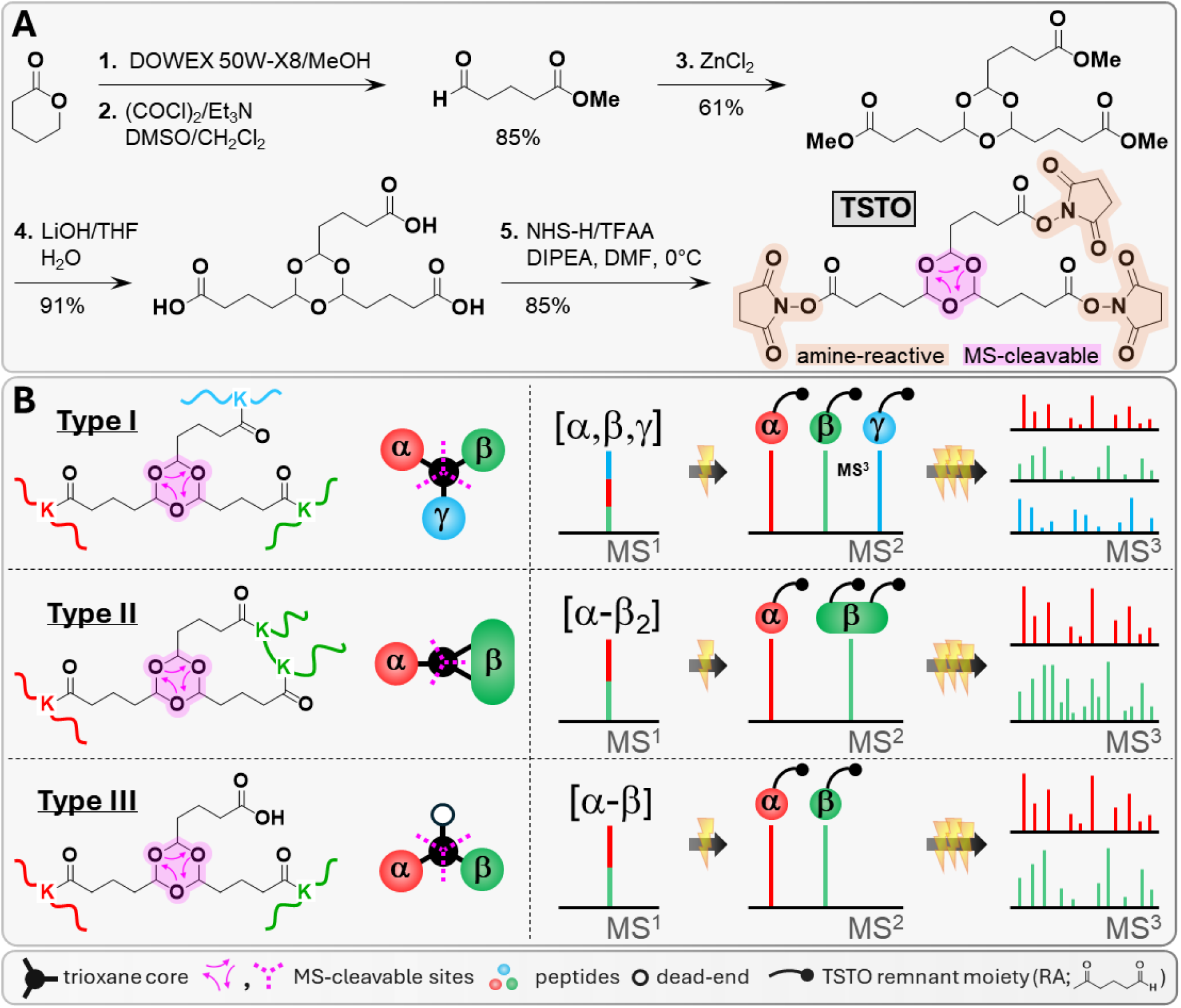
TSTO design, synthesis, and expected cross-link formation and MS^n^ fragmentation. **(A)** TSTO synthesis. **(B)** TSTO cross-link products exhibit unique fragmentation during CID analysis. For tripeptide tri-links (<u>Type I</u>), MS^2^ yields three fragment peptides with single AR-modified lysines. Dipeptide tri-links (<u>Type II</u>) produce two fragment peptides, one with a single AR-modified lysine and one with two. Finally, dipeptide bi-links (<u>Type III</u>) result in two fragment peptides, each with a single AR-modified lysine. MS^3^ analyses of these fragments enable unambiguous identification of TSTO tri- and di-links.

### Characterization of TSTO Cross-links

To characterize TSTO cross-linking and fragmentation, we first performed XL-MS analysis of the human 26S proteasome. Cross-linked proteasomes were digested and analyzed by LC MS^n^. As a result, the three types (I-III) of TSTO cross-linked peptides were identified and all displayed unique MS^2^ fragmentation as expected. This is illustrated by MS^n^ analyses of representative TSTO cross-linked peptides of the 26S proteasome (Figure S1). For a tripeptide tri-link [α, β, γ] (*m/z* 998.9354^5+^), cleavage of the trioxane using low-energy CID during MS^2^ yielded a series of dominant ions corresponding to α_AR_, β_AR_, and γ_AR_ peptides (Figure S1A). Notably, we observed that the aldehyde remnant can undergo water loss, leading to the detection of fragment ion pairs (i.e. AR and AR*). In addition to confirming the cross-link fragments, the mass difference between the ion pair (Δ=18.02 Da) can be used to selectively target TSTO cross-link fragment peptides for MS^3^ analysis, similar to the mass difference-based acquisition strategies used for sulfoxide-containing MS-cleavable cross-linked peptides ^1^. As illustrated (Figure S1B), MS^3^ analyses of the three cross-link fragments α_AR*_, β_AR*_, and γ_AR*_ enabled the identification of a tripeptide TSTO tri-link among Rpt1:K116, Rpt4:K206, and Rpt3:K238. For a bipeptide tri-link [α-β_2_] (*m/z* 1209.0961^4+^), MS^2^ fragmentation resulted in two sets of dominant ion species: α_AR_/α_AR*_, and β_2AR_/β_AR_AR*_/β_2AR*_ (Figure S1C). The detection of a fragment triplet with 18 Da increments indicates that peptide β carries two modified lysines, whereas peptide α only contains a single modified lysine. MS^3^ analyses of α_AR*_ and β_2AR*_ identified their sequences as ^278^AYEK_AR*_ILFTEATR^289^ and ^444^DGVIEASINHEK_AR*_GYVQSK_AR*_EMIDIYSTR^470^, respectively (Figure S1D), signifying a bipeptide tri-link [Rpn12:K281-Rpn3:K455, K461]. Finally, for a bipeptide bi-link [α-β] (*m/z* 773.4180^4+^), MS^2^ fragmentation resulted in two dominant ion pairs: α_AR_/α_AR*_, and β_AR_/β_AR*_ (Figure S1E); MS^3^ analyses of α_AR*_ and β_AR*_ determined a cross-link between Rpt4:K72 and Rpt6:K222 (Figure S1F). Owing to their unique fragmentation characteristics, TSTO cross-linked peptides were successfully identified by LC-MS^n^ using both intensity-(top) and mass difference-based (targeted) data-dependent acquisitions without requiring real-time precursor selection.

### TSTO XL-MS Analysis of the 26S Proteasome

In this work, the TSTO-based XL-MS analysis of the 26S proteasome has resulted in a total of 808 unique CSMs (cross-linked peptide spectra matches) (Table S1). Of these, 41.3% were tri-links, indicating that TSTO can efficiently cross-link three spatially proximal residues within the proteasome. To assess the accuracy of TSTO cross-links, we mapped them to a high-resolution 26S proteasome structure (PDB: 7QY7). Considering the spacer arm length of TSTO (∼14 Å), lysine residues with Cα-Cα distance ≤35 Å were expected to be preferentially cross-linked. As a result, all of the mapped tri-links and bi-links have ∼90% satisfaction rates (≤35 Å), supporting their validity (Figure S2). To understand whether the introduction of a supplementary reactive group is beneficial for describing protein topologies, we constructed a TSTO XL-PPI map of the 26S proteasome comprising 92 edges (Figure 2A). Compared to previous XL-MS studies using bifunctional linkers ^20–21^, 31 additional inter-subunit PPIs were identified—with most of them at interaction interfaces between proteasomal subcomplexes. To better understand the benefits of TSTO, we next examined the interactions captured by its tri-links. Among the 35 trimeric interactions within the proteasome, 13 described multimeric interactions among the six ATPase subunits (Rpt1-6), illustrating their proximity within the 19S base subcomplex (Figure 2B). In addition, the connectivity between the 19S and 20S subcomplexes was described by three trimeric interactions, including one among Rpt4, α1 and α7, and one between Rpt1 and α4 (Figure 2C,D). Apart from trimeric interactions involving three proteins, TSTO was able to concurrently place two distant residues of one protein (α4: K27, K166) to another (Rpt1:K418), due to their proximity in the three-dimensional structure. Another tri-link involving Rpt6 and α3 exemplifies the capability of TSTO to capture interactions involving adjacent residues (Rpt6:K397,K402) that may be missed by bifunctional linkers due to the tendency to form intra-links (loop-links). Moreover, TSTO cross-links were able to place a small proteasome subunit Dss1 in proximity to Rpn3, Rpn6 and Rpn7 within the 19S lid (Figures 2E, S3). Due to its small size (∼8 kDa) and disordered regions, Dss1 has not been easily resolved in high-resolution structures of the 26S proteasome until very recently ^8^. Although its interaction with Rpn3 and Rpn7 in the yeast proteasome has been determined by DSSO XL-MS ^22^, its localization had not been defined in the human 26S proteasome. Here, TSTO cross-linking confirmed Dss1 interactions with Rpn3 and Rpn7 in human proteasomes and also identified its contact with Rpn6 (Figure S3), localizing its interaction to the apical surface of the 19S lid. When mapped to the high-resolution structure, all proteasome-DSS1 cross-links corresponded to Cα-Cα distances below 35 Å with an average distance of 25 Å (Figure 2E, S3), further demonstrating the effectiveness of TSTO in triangulating protein interactions with precision. Collectively, these results have demonstrated that TSTO is effective in capturing multimeric interactions of proteasome complexes, providing three-dimensional contacts for the first time to support the spatial organization of these proximal subunits.

**Figure 2.**
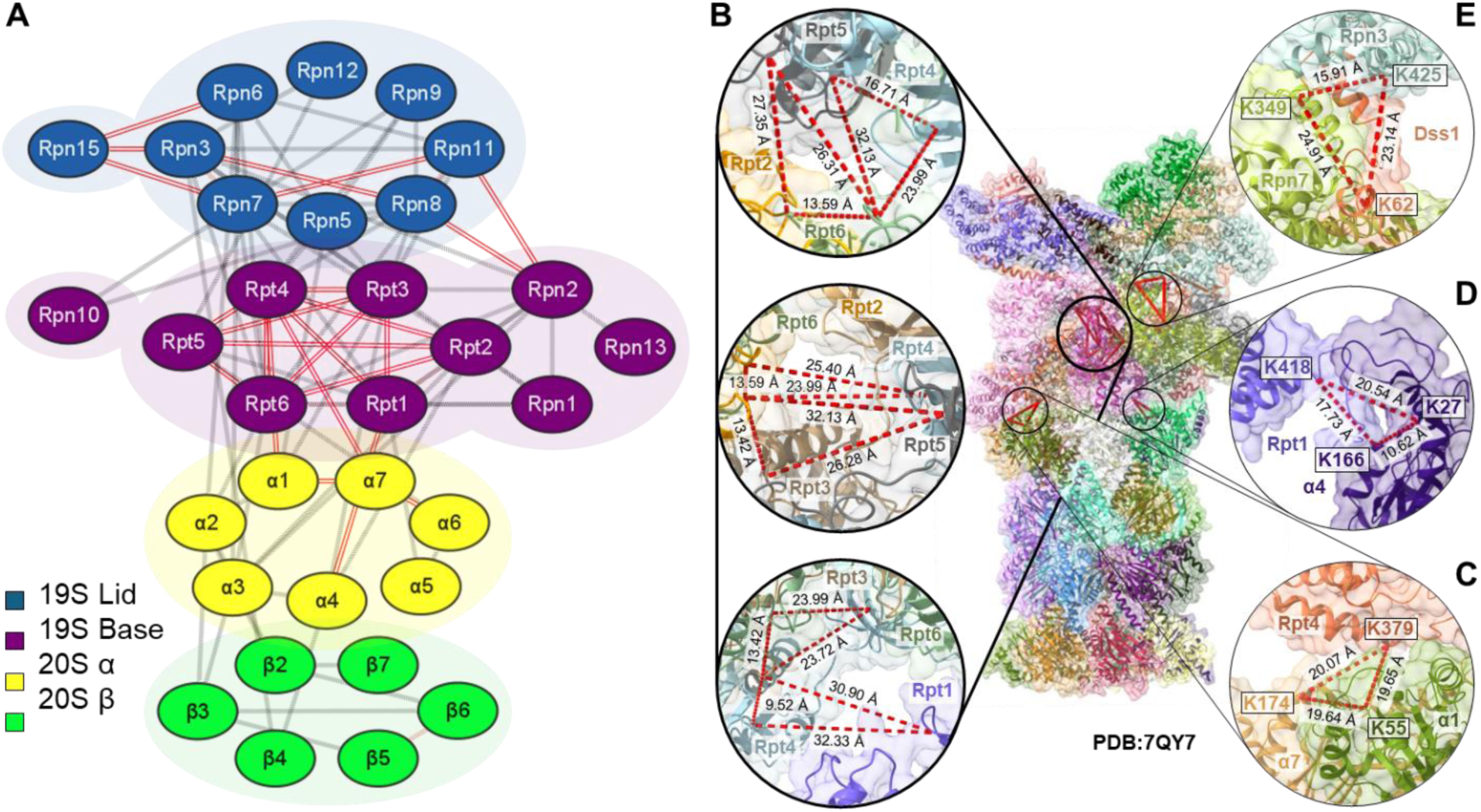
TSTO cross-linking of human 26S proteasome. **(A)** XL-PPI map of 26S cross-links. In total, 32 subunits were identified participating in inter-subunit interactions—19 from the 19S subcomplexes and 13 from the 20S subcomplex. 19S lid subunits shown in blue, 19S base subunits in purple, 20S α ring subunits in yellow, and 20S β ring subunits shown in green. Red lines represent interactions that have been captured as tri-links; grey lines represent other cross-links. Mapping of trimeric interactions to a high-resolution 26S proteasome structure: **(B)** lining the inner pore between Rpt subunits of the AAA-ATPase ring (19S base subcomplex), **(C, D)** Interactions describing connectivity of 19S base subcomplex subunits to the 20S, and **(E)** Dss1 with 19S lid subunits Rpn3 and Rpn7.

### In Vivo TSTO Cross-linking of the HEK293 Cells

We next investigated whether TSTO was applicable for system-wide XL-MS analysis to delineate cellular networks. Immunoblotting analysis indicated that TSTO is membrane permeable and suited for in-cell cross-linking (Figure S4). The general TSTO-based *in vivo* XL-MS workflow is illustrated in Figure 3, in which cross-linked peptides were subjected to two-dimensional peptide separation prior to LC-MS^n^ analysis ^23^. Across two biological replicates, we identified a total of 9079 unique CSMs (Table S2), of which 32.3% were tri-links. As shown, TSTO tri-links increased the total PPI yield by ∼27% compared to bi-link data alone (Figure S5A). Altogether, TSTO in-cell cross-linking yielded an XL-proteome of 1512 proteins containing 1242 PPIs (Figure S5B). These results indicate that TSTO cross-linking of intact cells is effective, and the presence of tri-links remains abundant in increasingly complex systems. Importantly, accurate identification of TSTO cross-links at the systems level was achieved using LC-MS^n^.

**Figure 3.**
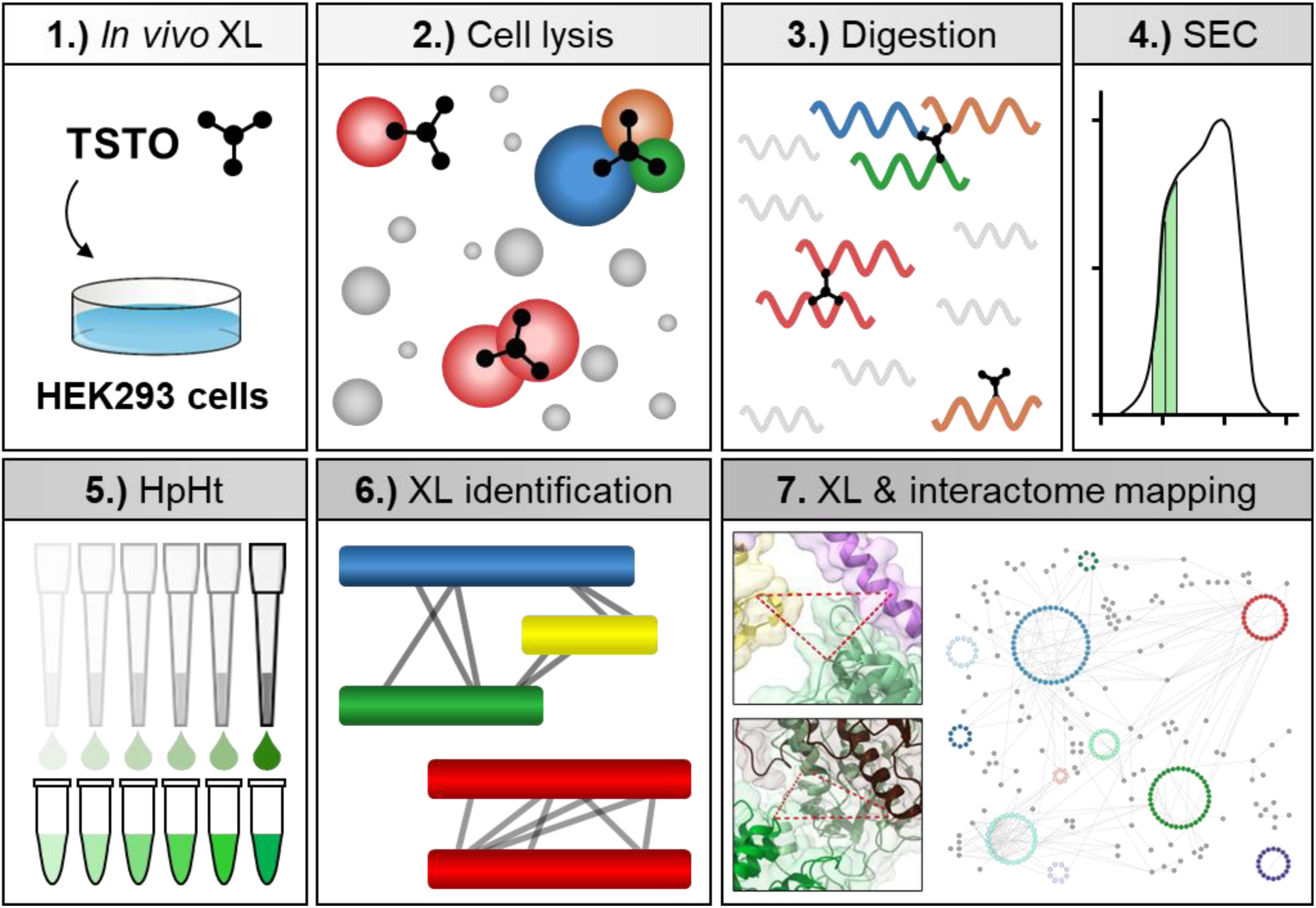
*In vivo* TSTO cross-linking workflow.

To examine the validity of the TSTO cross-links, we first mapped them to available high-resolution structures of protein complexes identified here (Figure S5C) ^9^. In total, 1790 K-K linkages were mapped across 539 CORUM complexes and 95% of them were considered satisfactory (≤ 35 Å). Next, we performed gene ontology (GO) analysis and confirmed that the TSTO XL-proteome covers a wide range of molecular functions, biological processes, and cellular components (Figure S6A). Compared to BioGRID and BioPlex databases, 48% of the TSTO XL-PPIs were known and 52% were novel. The STRING scores were found for 501 of the XL-PPIs and ∼70% were determined to be above 0.8 (Figure S6B), indicating high-confidence interactions ^9, 23^. Overall, among the 1242 inter-protein PPIs identified within this TSTO *in vivo* dataset, roughly one-third were novel when compared to an aggregate of recent systems-level cross-linking studies^23–27^. This suggests that while most XL-PPIs are supported, TSTO provides novel information not captured by other bifunctional MS-cleavable reagents. Collectively, these results demonstrate that TSTO has been successfully applied for global profiling of endogenous PPIs from their cellular environments.

### Multimeric Interactions of In Vivo Protein Complexes

Importantly, TSTO XL-MS has enabled the identification of multimeric interactions within various protein complexes. One well-represented complex is the 80S ribosome machinery. Specifically, extensive interactions describing subunit proximities were identified, particularly those between the 40S or 60S subcomplexes. This allows for an in-depth description of ribosomal PPIs with 3-D contacts (Figure S7). Interestingly, trimeric interactions were also found between 80S and several putative ribosome-binding partners, providing structural insights underlying their functional relevance in protein synthesis. Of these, the most frequent was SERBP1 (SERPINE mRNA-binding protein 1), which was shown to interact with 60S ribosome subunits RPL7A, RPL27, and RPL34 through several tri-links. While eukaryotic SERBP1 (Stm1 in yeast) has been associated with dormant ribosomes due to its role in clamping 40S and 60S subunits together to prevent mRNA access, it has been shown to primarily contact 40S subunits ^28–30^. Interestingly, TSTO identified ribosome-SERBP1 cross-links spanning the 80S, including distant 60S proteins.

In current high-resolution structures of ribosome-bound SERBP1, the majority of SERBP1 is unresolved—likely buried within the 80S. Together, the results suggest that the interaction of SERBP1 with ribosomal proteins is extensive, contacting various 40S and 60S subunits to inactivate 80S machinery. Another trimeric cross-link was identified between 40S ribosome subunits and UBAP2L (Ubiquitin-associated protein 2-like). While UBAP2L is a known RNA-binding protein (RBP) that may associate with ribosomes to facilitate protein synthesis ^31^, its function and roles remain poorly characterized. TSTO identified a trimeric interaction between UBAP2L and neighboring 40S subunits RPS7 and RPS27, triangulating its position near the small ribosomal complex and correlating well with the role of the 40S in initial binding and reading of mRNA.

One notable aspect is TSTO’s unique ability to define trimers, especially homomeric trimers (Table S2). This has been challenging for bifunctional linkers due to the difficulty in identifying multimeric interactions. For instance, the trimer of nucleoside-disphosphate kinase B (NME2) was detected due to the identification of a TSTO tri-link connecting three identical sequences containing NME2:K100. Interestingly, NME2 is known to form homohexameric structures comprising two stacked homotrimers. When mapped to a high-resolution structure of NME2 (PDB:8PYW), the loop regions containing the K100 residues of each homotrimer were found to localize along the axis of the hexamer, within 9.5 Å of one another (Figure 4A) ^32^. Similarly, homotrimeric cross-links were identified for the 10 kDa mitochondrial heat shock protein (HSPE1) through its K54 and K56 residues, suggesting an oligomeric complex. Indeed, HSPE1 has been shown to form a homoheptameric ring within the human mitochondrial chaperonin ’football’ complex (PDB: 4PJ1) ^33^, in which the longest distance spanned between any pair of HSPE:K54 or HSPE:K56 residues within neighboring triplets was 20 Å (Figure 4B). In addition, homotrimeric TSTO cross-links were identified from proteins that are known to assemble into oligomeric complexes but currently lack high-resolution structures. For instance, ATPase family AAA domain-containing protein 3A (ATAD3A) was shown to oligomerize through a specific residue (K262). Using AlphaFold3, we predicted the structure of an ATAD3A trimer at high confidence (90 > plDDT > 70) and mapped all possible K262 interactions satisfactorily (< 18.5 Å) (Figure 4C), exemplifying the capability of TSTO to facilitate structural modeling of protein oligomers.

**Figure 4.**
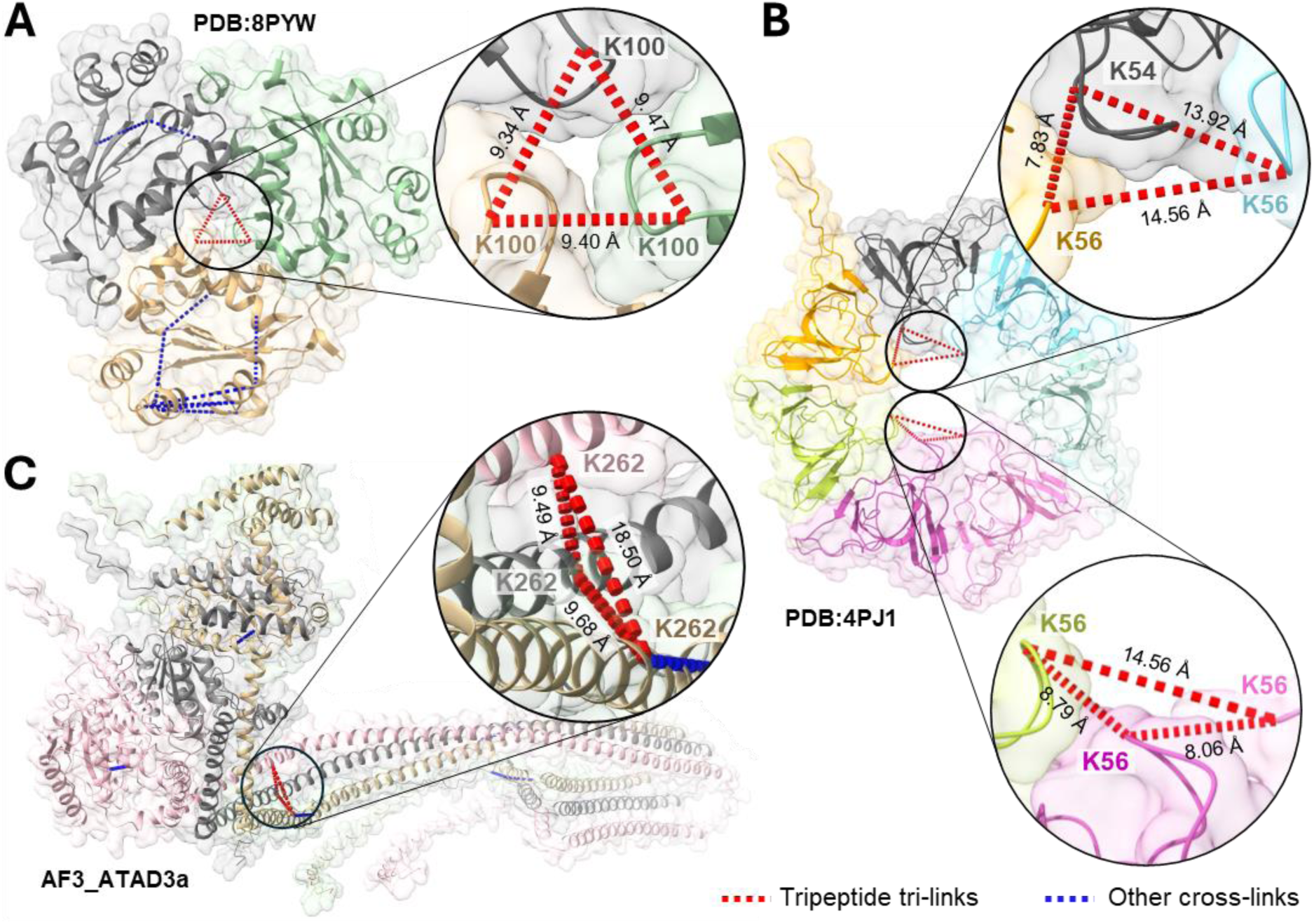
Mapping of homotrimeric TSTO cross-links from *in vivo* XL-MS. **(A)** Tri-link through K100 residues of three NME2 proteins, mapped to a high-resolution structure (PDB: 8PYW). **(B)** Trimeric cross-links involving residues K54 and K56 of three HSPE1 proteins mapped to a high-resolution structure (PDB:4PJ1). **(C)** Tri-link through K262 of three ATAD3a proteins, mapped to an AlphaFold3-generated model for an ATAD3a trimer.

## DISCUSSION

In this work, we have designed, synthesized, and characterized a novel trioxane-based, MS-cleavable, membrane-permeable homotrifuctional cross-linker, TSTO, to dissect multimeric protein interactions. This cross-linker enables simultaneous cross-linking of up to three proteins, allowing for more in-depth PPI analysis and providing additional restraints to advance XL-MS based structural analysis of protein assemblies. We have demonstrated that all types of TSTO cross-linked peptides display unique and predictable CID-induced fragmentation and can be unambiguously identified using LC MS^n^ analysis. TSTO’s unique capability to concurrently cleave all three cross-link arms and leave an identical remnant on each cross-linked residue establishes it a brand-new class of MS-cleavable reagent. Additionally, this distinctive feature minimizes the number of theoretical MS fragments corresponding to each peptide constituent, simplifying ion selection for subsequent MS^3^ analysis. While MS^2^-based data acquisitions have become popular for analyzing MS-cleavable bi-linked peptides, we envision that MS^3^-based approaches will be preferred for the identification of tripeptide cross-links. The co-fragmentation of three peptides within a single spectrum heavily convolutes database searching, impeding identification of higher-order cross-linked species by MS^2^-based approaches. Moreover, the central trioxane presents a novel core structure for developing multifunctional MS-cleavable cross-linkers as one arm can be replaced with other functional groups—such as enrichment, isobaric, or reporter tags—to enable cross-link purification, quantitation, or further improve cross-link detection and identification. The NHS ester groups within TSTO can also be replaced by other reactive chemistries to target specific or non-specific amino acids ^24, 34–36^, expanding the coverage of XL proteomes. As with TSTO, the cleavage of the trioxane within the mass spectrometer would release any additional functional groups, preventing their impact on cross-linked peptide identification. Thus, the development of TSTO presented here opens a new direction for designing diverse cross-linkers to further advance XL-MS technologies.

Compared to cross-linking reagents that target two residues, TSTO enhances the localization of interacting proteins by simultaneously targeting three residues, triangulating a multi-point attachment that provides greater spatial resolution and allows more precise mapping of protein interfaces and interaction sites. This is particularly useful for modeling multimeric interactions, especially for protein complexes that can exist in different compositional states. As bifunctional cross-linkers can only provide spatial restraints between two residues, it can be difficult to correctly assign them to individual compositional forms without reiterative modeling. Therefore, trimeric cross-links offer an additional restraint to help facilitate characterization of multi-protein interactions. As shown with cross-linking of the 26S proteasome, TSTO has identified trimeric cross-links bridging subunits of different proteasome subcomplexes that accurately describe their positioning within the 26S holocomplex—such as those between the 19S lid and base, and the 19S base and 20S subcomplexes. In addition, these trimeric links have also accurately described the orientation of multi-subunit interactions within their respective subcomplexes, such as the localization of Dss1 to the outer 19S lid and the close proximities of solvent-accessible residues of the ATPase ring subunits lining the central pore of the 19S base.

Similarly, TSTO has identified trimeric interactions from *in vivo* XL-MS, for instance triangulating the positions of ribosome-interacting proteins relative to the larger 80S ribosomal machinery, as well as various subunit interfaces within and between its 40S and 60S subcomplexes. In addition to revealing the identities of heteromeric interactions, TSTO expands the capability of XL-MS to identify homomeric ones. While binary cross-linkers can reveal some homodimer formation by identifying cross-linked peptides with overlapping sequences, TSTO is capable of placing homodimeric interactions in the three-dimensional context of multiprotein assemblies by reacting with a third residue, providing a clearer picture of how individual dimers are oriented as part of a larger protein complex. Furthermore, TSTO can identify homotrimeric cross-links—as confirmed for NME2 and HSPE1—which can be used in conjunction with structure prediction such as AlphaFold to model unknown structures.

In summary, we have successfully established and employed the TSTO-based XL-MS platform to capture trimeric interactions of protein complexes and cellular networks from intact cells, providing new structural details that cannot be easily obtained by existing reagents. The information obtained can be combined with AlphaFold prediction and/or integrative structural modeling to enhance the characterization of cellular networks in future studies. Therefore, the TSTO-based XL-MS platform represents a highly promising approach for advancing XL-MS technology towards systems structural biology *in vivo*.

## MATERIALS AND METHODS

### TSTO Synthesis

Starting with delta-valerolactone, (1) acid-catalyzed ring opening in methanol provided a methyl ester, which was (2) converted to an aldehyde via Swern oxidation in an 85% yield over two steps. (3) The resulting aldehyde was treated with 10 mol% zinc-(II) chloride, affording a symmetric trioxane tris-ester. (4) Hydrolysis of the tris-ester afforded a tris-acid, which was (5) converted to TSTO by ester formation with NHS and TFAA.

### Affinity Purification and TSTO Cross-linking of Human 26S Proteasomes

A stable 293^HBTH-Rpn1^ cell line ^37^ was first grown to confluency. After native cell lysis, human 26S proteasomes were purified from the clarified lysate by binding to streptavidin– sepharose resin. Bead-bound proteasomes were cross-linked on-bead in PBS buffer (pH 7.5) with 0.75 mM TSTO for 1 hr at room temperature. After quenching the cross-linking reaction using ammonium bicarbonate, the proteins were reduced with tris(2-carboxyethyl)phosphine (TCEP) for 30 min at RT and alkylated with iodoacetamide in the dark at RT for 30 min. Cross-linked proteins were then digested in 8M urea buffer using LysC for 4 hrs at 37 °C, followed by trypsin digestion at 37 °C overnight after diluting urea concentration to < 1.5 M. The resulting peptide mixtures were extracted and desalted using C18 tips (Agilent) prior to MS analyses.

### Optimization of In Vivo TSTO Cross-linking

A stable 293^HBTH-CSN2^ cell line ^38^ was first grown to confluency, washed with PBS, and then gently pelleted. To determine the optimal cross-linking conditions, intact cells were cross-linked at various cross-linking concentrations ranging from 0.5 to 3 mM, at room temperature or 37 °C. Clarified lysates from each condition were separated by SDS-PAGE and transferred onto a PVDF membrane and stained using amido black. After rinsing off the dye and blocking using 5% milk in TBST, the membrane-bound proteins were incubated with streptavidin-HRP to monitor the oligomization of HBTH-CSN2 in response to cross-linking conditions. Based on these results, *in vivo* cross-linking of intact HEK293 cells was performed at 1 mM.

### TSTO Cross-linking of Intact Cells

Intact 293^HTBH-CSN2^ cells were cross-linked using 1 mM TSTO in PBS buffer (pH 7.4) for 1 hr with rotation at room temperature. Afterwards, the cross-linking reaction was quenched using excess ammonium bicarbonate (50 mM) for 10 min. Cells were spun down, washed again with PBS, and needle-lysed on ice in denaturing buffer (8 M urea, 50 mM Tris-HCl pH 7.0). Lysate was clarified by centrifugation at 21,000g for 13 min in 4 °C and the resulting supernatant was transferred to 30,000 NMWL Microcon centrifugal tubes for FASP digestion similarly as described ^39^. Briefly, proteins atop the filter were reduced using 10 mM TCEP for 20 min at RT, alkylated using 20 mM iodoacetamide in the dark at RT for 20 min, and then digested in 8M urea buffer using LysC for 4 hrs at 37°C followed by trypsin digestion at 37 °C overnight after urea dilution. The resulting cross-linked peptide mixtures were then spun through the filter and desalted using Waters C18 Sep-Pak cartridges.

### SEC-HpHt Enrichment of Cross-linked Peptides

Peptide separation by SEC was performed similarly as described ^40^. Briefly, dried peptides were reconstituted in SEC mobile phase (0.1% formic acid and 30% ACN) and separated on a Superdex Peptide PC 3.2/30 column (300 x 3.2 mm) at a flow rate of 50 µL/min, monitored at 215, 254 and 280 nm UV absorbance. Two-minute fractions were collected, and only fractions 24 and 26 containing the most cross-linked peptides were collected. For high-pH separation, HpHt tips were prepared as described ^23^, and SEC-separated fractions were vacuum dried and resuspended in ammonium water (pH 10). After loading onto the tip, the peptides were eluted using increasing percentage of ACN in ammonia water (6%, 9%, 12%, 15%, 18%, 21%, 25%, 30%, 35%, and 50%). The 25%, 30%, 35% and 50% fractions were then combined with 6%, 9%, 12% and 21% fractions, respectively. The final SEC-HpHt fractions were vacuum dried and stored at -80 oC before MS analysis.

### LC-MS^n^ Analysis

LC-MS^n^ analysis of cross-linked peptides was performed using an UltiMate 3000 UPLC (Thermo Fisher Scientific) liquid chromatograph coupled on-line to an Orbitrap Fusion Lumos mass spectrometer (Thermo Fisher Scientific). Peptides were separated by reverse-phase on a 50cm x 75μm I.D. Acclaim® PepMap RSLC column using gradients of 4% to 25% acetonitrile at a flow rate of 300 nL/min (solvent A: 100% H_2_O, 0.1% formic acid; solvent B: 100% acetonitrile, 0.1% formic acid) prior to MS^n^ analysis. For each MS^n^ acquisition, duty cycles consisted of one full Fourier transform scan mass spectrum (375–1500 m/z, resolution of 60,000 at *m/z* 400) followed by data-dependent MS^2^ and MS^3^ acquired at top speed in the Orbitrap and linear ion trap, respectively. Ions detected in MS^1^ with 4+ or greater charge were selected and subjected to CID fragmentation (NCE 23%) in MS^2^ and resulting ions were detected in the Orbitrap (resolution 30,000). Ions observed in MS^2^ spectra with charge 2+ or greater were selected and fragmented in MS^3^ using CID (NCE 35%) and detected in the linear ion trap in ‘Rapid’ mode. Ions selected for MS^3^ were either based on abundance (top 4 or 5 most intense ions in MS^2^) or targeted based on doublets with mass difference pairs (Δ=18.02 Da) corresponding to cross-linker remnant moiety water loss. For 26S proteasome cross-links, each acquisition was 200 min and ion selection for MS^3^ was based on top intensity. For SEC-HpHt *in vivo* samples. each acquisition was 120 or 150 min, and both top intensity and mass difference-targeted methods were used for selecting ions for MS^3^ analysis.

### Identification of TSTO Cross-links by MS^n^

Spectrometric data were extracted from .raw files using PAVA ^41^. Extracted MS^3^ spectra were subjected to protein database searching via Batch-Tag within a developmental version of Protein Prospector (v. 6.3.5, University of California, San Francisco) against a randomly concatenated SwissProt database consisting of 20,418 human proteins and their corresponding decoys. Mass tolerances for parent ions and fragment ions were set as ± 15 ppm and 0.6 Da, respectively. Trypsin was set as the enzyme with three maximum missed cleavages allowed. Cysteine carbamidomethylation was selected as a constant modification, while protein N-terminal acetylation, methionine oxidation, and N-terminal conversion of glutamine to pyroglutamic acid were selected as variable modifications. Two additional defined variable modifications on uncleaved lysines and free protein N-termini were selected: AR (C_5_H_6_O_2_, 98.04 Da) and AR* (C_5_H_4_O, 80.03 Da), corresponding to remnant moieties for cleaved TSTO. MS^n^ data (monoisotopic masses and charges of parent ions and corresponding fragment ions and MS^3^ database search results were integrated via in-house software xl-Tools ^42^ to automatically generate, summarize and validate identified cross-linked peptide pairs. Experimental FDR calculated using a target-decoy approach was determined to be 0.01% (3/28,949) at the CSM level.

## ACKNOWLEDGMENTS

We wish to thank Prof. A.L. Burlingame, Drs. Peter Baker, and Robert Chalkley for their support of Protein Prospector, and the Huang lab members, especially Paul Morenkov, for their help. This work was supported by National Institutes of Health grants R35GM145249 to L.H., and R35GM151256 to I.E..

## AUTHOR CONTRIBUTIONS

L.H. conceived the study and directed the research. C.Y., X.W., E.N, S.R. and L.H. designed the experiments. C.Y. performed XL-MS experiments, data acquisition and analyses. X.W. purified and cross-linked proteasome complexes, E.N. and S.R. performed cross-linker synthesis, I.E. performed alpha fold modeling. C.Y. and L.H. wrote the manuscript with contributions from other authors.

