## Supplementary Information for "Trioxane-based MS-cleavable Cross-linking Mass Spectrometry for Profiling Multimeric Interactions of Cellular Networks"

Medical Science I, D233

Department of Physiology & Biophysics

University of California, Irvine

Irvine, CA 92697-4560

**TABLE OF CONTENTS**

Supplementary Tables ... 3

Supplementary Figures ... 4

Figure S1 ... 4

Figure S2 ... 6

Figure S3 ... 7

Figure S4 ... 8

Figure S5 ... 9

Figure S6 ... 10

Figure S7 ... 11

References ... 12

#### Supplementary Tables

**Table S1.** TSTO cross-links identified from affinity-purified human 26S proteasomes.

**Table S2.** TSTO cross-links identified from *in vivo* cross-linked HEK 293 cells.

### Supplementary Figures

#### Figure S1

**Representative LC-MS$^n$ analyses of the three types of TSTO cross-links identified from 26S proteasomes. (A)** MS$^2$ fragmentation of a tripeptide tri-link [α, β, γ] with precursor *m/z* 998.9354$^{5+}$ yielded a series of dominant ion doublets corresponding to $\alpha_{AR}/\alpha_{AR*}$, $\beta_{AR}/\beta_{AR*}$, and $\gamma_{AR}/\gamma_{AR*}$ peptides. **(B)** MS$^3$ analyses of the cross-link fragments $\alpha_{AR*}$ (*m/z* 652.3762$^{2+}$), $\beta_{AR*}$ (*m/z* 895.4729$^{2+}$), and $\gamma_{AR*}$ (*m/z* 922.9833$^{2+}$) identified them as $^{110}$YIINVK$_{AR*}$QFAK$^{120}$, $^{198}$VVSSSIVDK$_{AR*}$YIGESAR$^{213}$, and $^{230}$VVGSEFVQK $_{AR*}$YLGEGPR$^{245}$, signifying a tripeptide tri-link among Rpt1:K116, Rpt4:K206, and Rpt3:K238. **(C)** MS$^2$ fragmentation of a bipeptide tri-link [α-$\beta_2$] (*m/z* 1209.0961$^{4+}$) yielded two dominant sets of ions—a doublet representing $\alpha_{AR*}/\alpha_{AR}$ and a triplet representing $\beta_{2AR}/\beta_{AR\_AR*}/\beta_{2AR*}$ peptides. **(D)** MS$^3$ analyses of the cross-link fragments $\alpha_{AR*}$ (*m/z* 761.4022$^{2+}$) and $\beta_{2AR*}$ (*m/z* 1086.8612$^{2+}$) identified their sequences as $^{278}$AYEK$_{AR*}$ILFTEATR$^{289}$ and $^{444}$DGVIEASINHEK$_{AR*}$GYVQSK$_{AR*}$EMIDIYSTR$^{470}$ respectively, signifying a dipeptide tri-link among Rpn12:K281, Rpn3:K455, and Rpn3:K461. **(E)** MS$^2$ fragmentation of a dipeptide bi-link [α-β] (*m/z* 773.4180$^{4+}$) resulted in two dominant ion doublets $\alpha_{AR}/\alpha_{AR*}$ and $\beta_{AR}/\beta_{AR*}$. **(F)** MS$^3$ analyses of the cross-link fragments $\alpha_{AR*}$ (*m/z* 591.8356$^{2+}$) and $\beta_{AR*}$ (*m/z* 878.9664$^{2+}$) identified their sequences as $^{69}$FIVK$_{AR*}$ATNGPR$^{78}$ and $^{214}$VSGSELVQK$_{AR*}$FIGEGAR$^{229}$, signifying a pair-wise cross-link between Rpt4:K72 and Rpt6:K222. Note: AR: aldehyde remnant moiety; AR*: aldehyde remnant moiety after water loss (i.e. AR-$H_2O$).

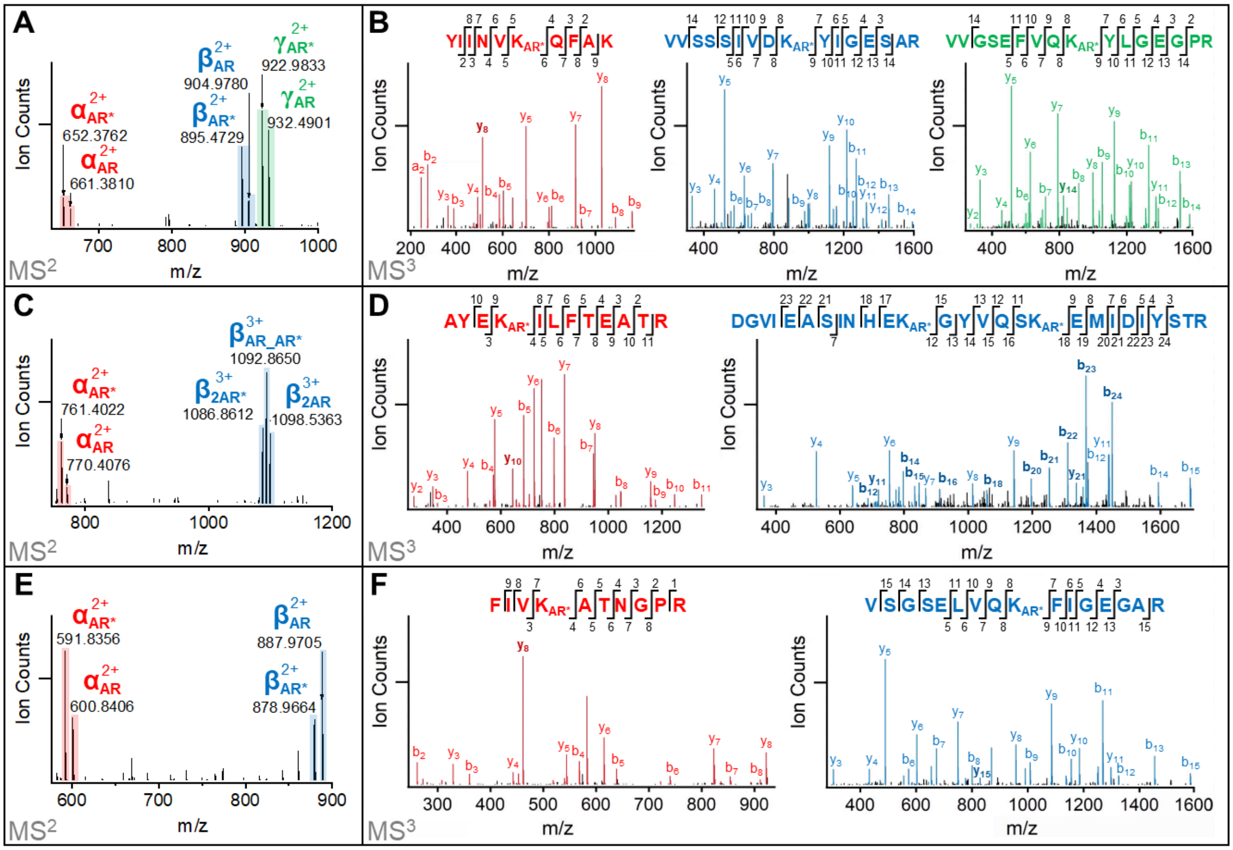
A
Ion Counts
α AR* 2+
652.3762
α AR 2+
661.3810
β AR 2+
904.9780
β AR* 2+
895.4729
γ AR* 2+
922.9833
γ AR 2+
932.4901
700 800 900 1000
m/z
MS2
B
YIINVKAR*QFAK
VVSSSIVDKAR*YIGESAR
VVGSEFVQKAR*YLGEGPR
Ion Counts
m/z
MS3
C
Ion Counts
α AR* 2+
761.4022
α AR 2+
770.4076
β AR_AR* 3+
1092.8650
β 2AR* 3+
1086.8612
β 2AR 3+
1098.5363
800 1000 1200
m/z
MS2
D
AYEKAR*ILFTEATR
DGVIEASINHEKAR*GYVQSKAR*EMIDIYSTR
Ion Counts
m/z
MS3
E
Ion Counts
α AR* 2+
591.8356
α AR 2+
600.8406
β AR 2+
887.9705
β AR* 2+
878.9664
600 700 800 900
m/z
MS2
F
FIVKAR*ATNGPR
VSGSELVQKAR*FIGEGAR
Ion Counts
m/z
MS3

**Figure S2**

**Distance distribution plots of unique residue pairs derived from TSTO cross-linking of 26S proteasomes**. Unique residue pairs from TSTO bi-links are shown in gray, while those from TSTO tri-links are shown in light blue. Cross-linked residues were mapped onto a high-resolution structure of the 26S proteasome (PDB: 7QY7).

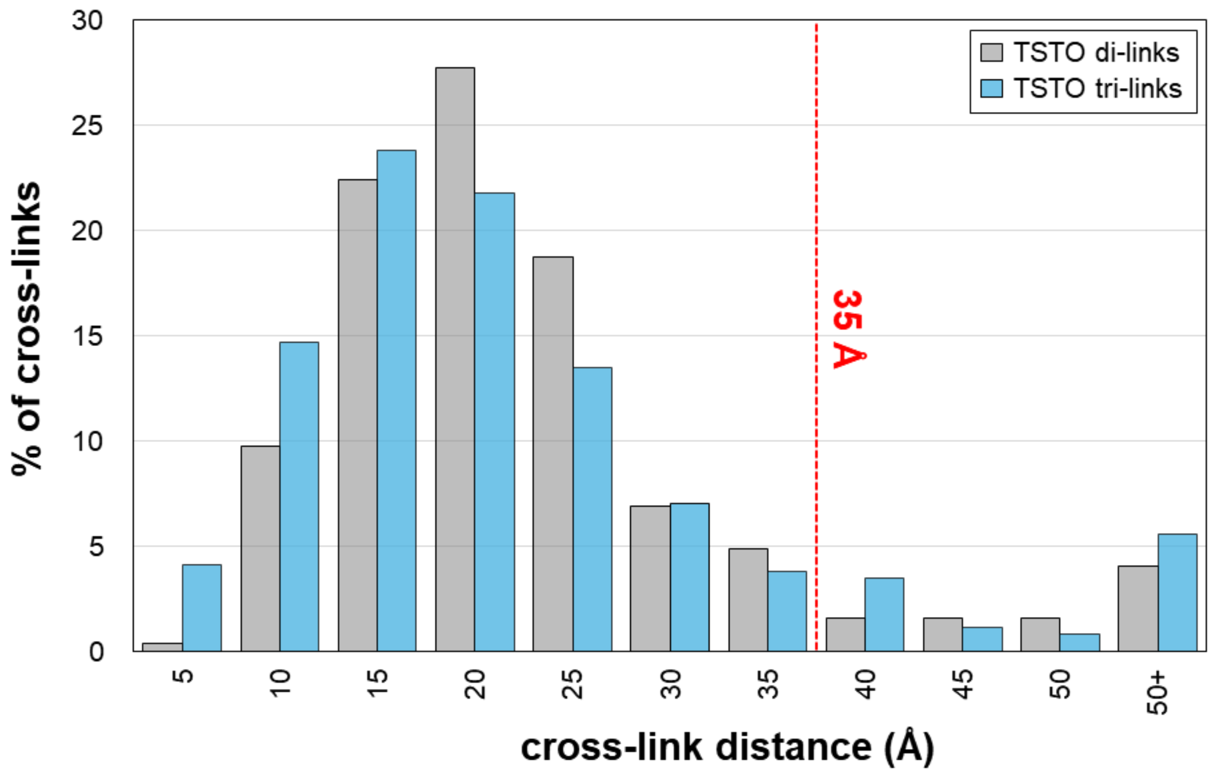

#### Figure S3

**Dss1 interactions captured by TSTO cross-linking. (A)** 2-D XL-map of Dss1 cross-links to 19S lid subunits. Trimeric interactions are highlighted in red. **(B)** Dss1 cross-links to 19S proteasome lid mapped to high-resolution 26S proteasome structure (PDB:7QX7).

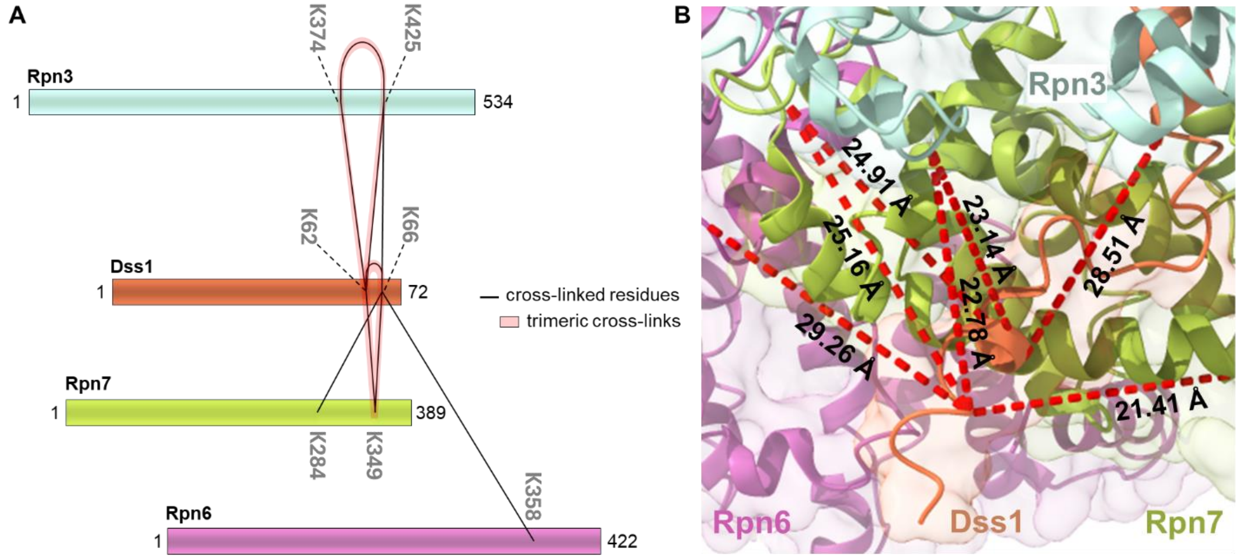

#### Figure S4

**Evaluation and optimization of TSTO *in vivo* cross-linking.** TSTO cross-linking was tested at various concentrations (0.5-3 mM) using HEK 293$^{HTBH\text{-}CNS2}$ cells. The cross-linked products were separated by SDS-PAGE, transferred onto a PVDF membrane, and evaluated by **(A)** amido black staining and **(B)** immunoblot analysis using StrepHRP to probe HTBH-tagged CSN2. Emphasized band corresponds to monomeric HTBH-tagged CSN2.

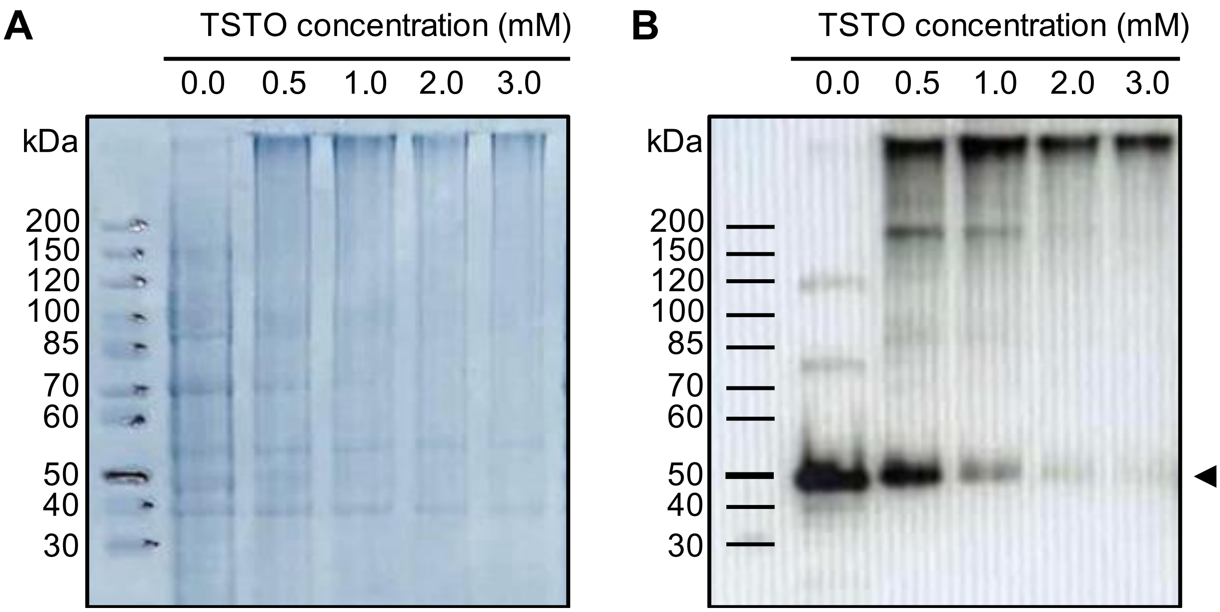

#### Figure S5

**26S TSTO cross-linking analysis. (A)** Venn diagrams depicting the overlap of inter- and intra-protein PPIs described by TSTO bi- and tri-links captured from *in vivo* cross-linking. **(B)** Histogram of mapped Ca-Ca distances for 1790 URPs across 539 CORUM complexes; 95% were found to be ≤ 35 Å. **(C)** *In vivo* XL-PPI network of HEK 293 cells derived from TSTO cross-links comprising 1512 nodes connected by 1242 edges.

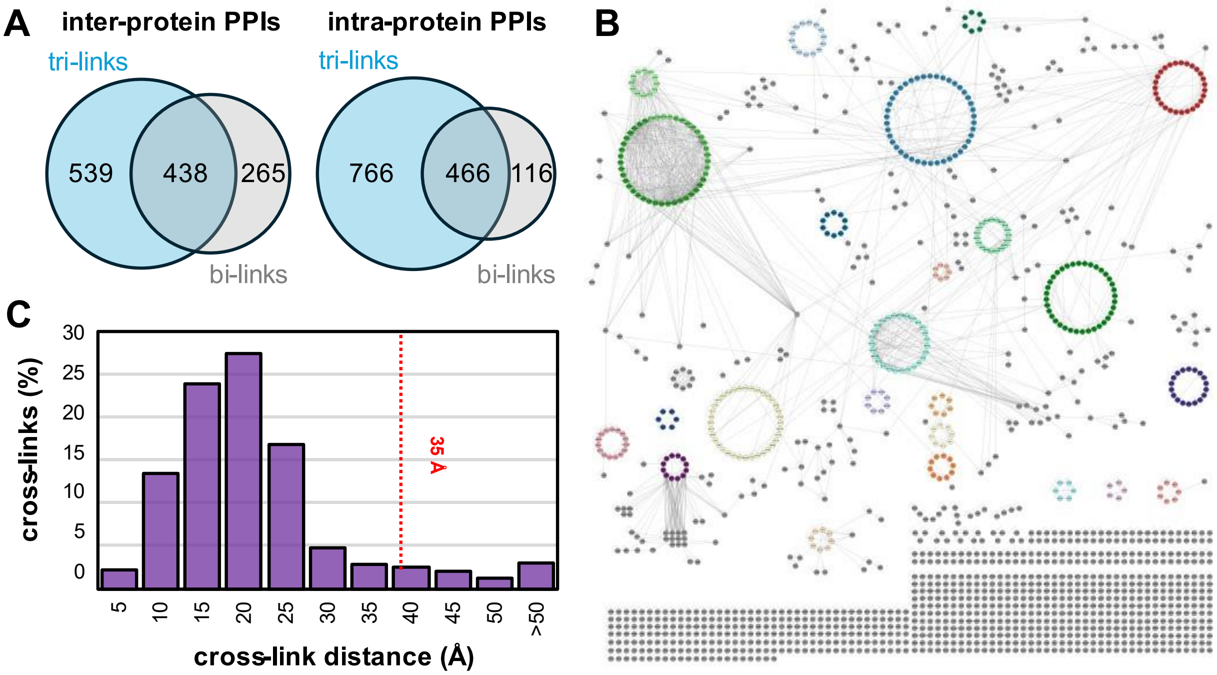

#### Figure S6

**Evaluation of the TSTO XL-Proteome. (A)** Gene Ontology (GO) analysis showing cell compartment distribution of TSTO XL-proteome compared to GO proteome and DSBSO XL-proteome [1]. In both cross-linking datasets, cytosolic proteins are enriched, and plasma membrane proteins have decreased representation compared to the Gene Ontology proteome. (**B)** Distribution of STRING scores for XL-PPIs captured by *in vivo* TSTO cross-linking compared to human PPIs curated within the STRING database.

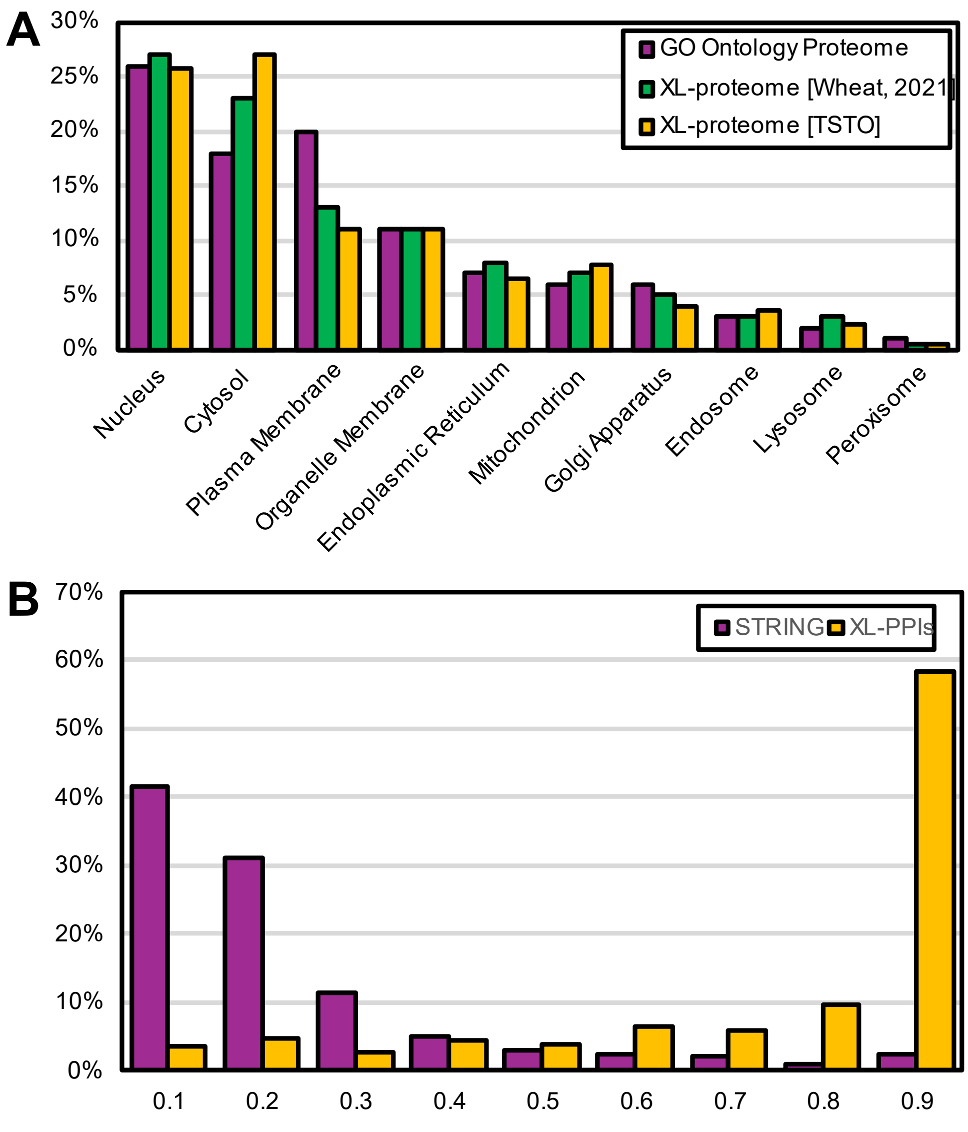

#### Figure S7

**Mapping of TSTO cross-links to a high-resolution structure of the 80S ribosomal complex (PDB: 6Z6M).** Mapped distances between residues identified in trimeric cross-links are shown in red, while other cross-links are shown in blue. The overall satisfaction rate of mapped cross-links ≤ 35 Å was 96%.

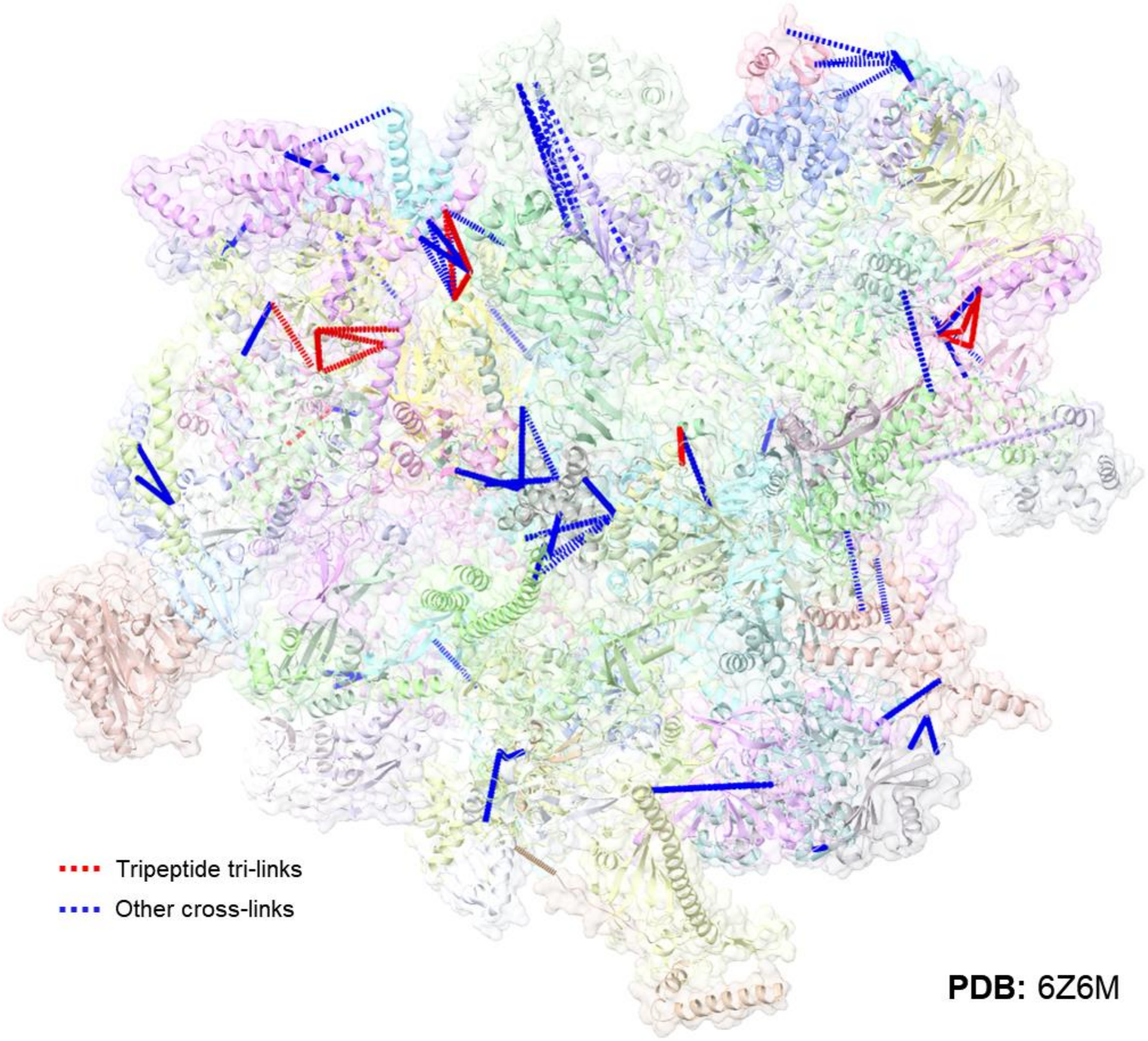
